# A neutralizing human antibody induces movement of the HCoV-229E receptor binding domain

**DOI:** 10.1101/2025.05.14.654097

**Authors:** Mera F. Liccione, Trevor Scobey, Heather M. Froggatt, Camryn Pajon, Boyd L. Yount, Qi Yin, Taylor N. Spence, Emmanuel B. Walter, Kevin O. Saunders, Robert J. Edwards, Ralph S. Baric, Barton F. Haynes, Derek W. Cain, Timothy P. Sheahan, Daniel Wrapp

## Abstract

HCoV-229E is an endemic *Alphacoronavirus* that typically causes common cold-like disease in most healthy adults, but can also cause severe respiratory disease in the very young and the elderly. Although the virus was discovered over sixty years ago and undergoes continuous antigenic drift, remarkably little is known about the humoral immune response to HCoV-229E infection. Here we report the isolation of two receptor binding domain-targeting neutralizing human antibodies raised in response to natural HCoV-229E infection. One of these, DH1533, potently neutralizes HCoV-229E, binds to spike with sub-nanomolar affinity and prevents the association between the RBD and the host cell receptor aminopeptidase N. Structural characterization of this antibody bound to HCoV-229E spike delineated a neutralization-sensitive epitope on the RBD and revealed that DH1533 induces conformational flexibility in neighboring RBDs, reminiscent of the “up-and-down” kinetics observed in the related *Betacoronavirus* spikes. These findings provide insight into the humoral immune response to HCoV-229E infection and will serve as a guide for the design of future therapeutic interventions.

## INTRODUCTION

To date, there are seven prominent human coronaviruses (HCoVs) which cause mild to severe respiratory infection. In the past 20 years, three novel CoVs have emerged from animal reservoirs resulting in endemic, epidemic and pandemic disease^1–3^. Beyond these seven pathogens, there is considerable potential for the future emergence of novel HCoVs as evidenced by the identification of multiple enzootic CoV in animal reservoirs capable of binding to and entering human cells^4–9^, as well as several CoVs that have already been shown to cause sporadic infections in human hosts^10–12^. While most HCoVs, including MERS-CoV and SARS-CoV-2, belong to the *Betacoronavirus* genus, two seasonal endemic HCoVs, HCoV-NL63 and HCoV-229E, belong to the *Alphacoronavirus* genus^13^. HCoV-229E was initially isolated from a human patient in 1962^14^ and is known to cause bronchitis and mild respiratory disease, although these symptoms can be more severe in infants, the elderly, and the immunocompromised^15^. Exposure to HCoV-229E is thought to be nearly universal by five years of age^16^, although surprisingly little is known about the immune response to this ubiquitous pathogen, its evolution in response to human herd immunity, and the factors that mediate the development of mild or severe disease.

Binding, fusion and entry into host cells by HCoV-229E is mediated by a heavily glycosylated class I viral fusion protein called spike (S)^17^. Like other CoV S, the homotrimer HCoV-229E S can be divided into two functionally distinct subunits, the S1 subunit which contains the receptor binding machinery^18^, and the S2 subunit which contains the hydrophobic fusion peptide and the alpha helical heptad repeats that drive membrane fusion^19^. Binding of the host cell receptor aminopeptidase N (APN) to the receptor binding domain (RBD) in the S1 subunit gradually destabilizes the prefusion conformation of S, ultimately leading to S1 dissociation and the irreversible conformational transition from the prefusion to the postfusion conformation^20–22^.

Although the *Alphacoronavirus* S proteins share many functional similarities with those from the more rigorously characterized *Betacoronavirus* genus, they also diverge in a number of key ways. For example, whereas the *Betacoronavirus* RBD rests on top of the N-terminal domain (NTD) of the neighboring protomer^23^, the HCoV-229E RBD points back in the opposite direction, such that its receptor binding loops interact with the NTD of the same protomer^17,24,25^. Additionally, many *Betacoronavirus* RBDs are known to stochastically sample an “up” conformation, in which a hinge-like movement fully exposes the RBD, allowing it to bind to the host cell receptor free of steric hindrances introduced by the rest of S^26,27^. While the bulky, globular structure of APN makes it clear that the HCoV-229E RBD must undergo an analogous conformational change^22^, it appears to sample this up conformation much less frequently than the *Betacoronavirus* RBDs. In fact, this conformation was only recently observed for the first time by cryo-EM, where it was captured by incubating APN and HCoV-229E S together to trap the RBD up^20^. Despite this major breakthrough, the molecular mechanisms that mediate this critical conformational change remain poorly understood.

In addition to these complex conformational dynamics, HCoV-229E S is also continuously undergoing antigenic drift, presumably as a means of evading neutralizing antibodies that have been elicited by prior HCoV-229E exposure^28^. Reinfection with HCoV-229E is thought to occur every ~2-4 years^29,30^, suggesting that in most cases, infection elicits a humoral immune response that then protects from subsequent challenges until the viral genetic drift is sufficient to evade prior immunity. This phenomenon has been elegantly investigated using historical patient sera samples and a variety of HCoV-229E pseudoviruses meant to recapitulate large swaths of antigenic diversity^28^; but without neutralizing monoclonal antibodies and high-resolution structural information about their binding epitopes, an understanding of the precise molecular determinants that govern this evolutionary arms race remains elusive.

Here we report the isolation of two neutralizing HCoV-229E RBD-directed monoclonal antibodies (mAbs) from an individual recovering from recent HCoV-229E infection. These antibodies bind with sub-nanomolar affinities, and one of them, DH1533, is capable of preventing the interaction between the HCoV-229E RBD and APN. By determining the cryo-EM structure of DH1533 bound to HCoV-229E S, we identified its mechanism of neutralization, providing a structural rationale for strain-specific viral evasion. Furthermore, our cryo-EM studies reveal that DH1533 binding induces movement of the RBDs in neighboring unbound protomers, potentially shedding light on the conformational dynamics that govern RBD accessibility. These studies provide novel insights into how the humoral immune system is able to target a neutralization-sensitive epitope on the surface of prefusion HCoV-229E S. These findings have important implications for the development of sorely needed therapeutic interventions for this CoV genus, which will be essential for future pandemic preparedness.

## RESULTS

An elderly individual (≥65 years of age) reporting mild respiratory disease tested positive for HCoV-229E infection by RT-PCR, although a clinical isolate of the infecting strain could not be obtained. Symptoms resolved over three days and peripheral blood mononuclear cells (PBMCs) were collected approximately two weeks after the onset of symptoms. Based on the age of the donor, it is highly unlikely that this was their first HCoV-229E infection, although confirmation of prior exposure could not be definitively determined. Antigen-specific B cells were isolated by fluorescence-activated cell sorting (FACS) using fluorochrome-conjugated HCoV-229E RBD-SD1 (residues 277-495 from isolate B1/GER/2015) tetramers as staining antigens (**Figure 1A, Supplementary Figure 1**). A total of 103 antibody sequences were determined from sorted B cells by nested PCR. Two of the corresponding cloned monoclonal antibodies (DH1532, DH1533) exhibited high-affinity binding to the HCoV-229E RBD-SD1 (**Figure 1B**), with DH1532 binding with an affinity of 205 pM and DH1533 binding with an affinity of 791 pM. DH1532 is derived from VH3-30*04/VK1-39*01 gene segments and DH1533 is derived from VH3-23*04/VK1-33*01 gene segments, but despite their high binding affinities, only DH1532 showed evidence of extensive somatic hypermutation (DH1532 V_H_=9.0%, DH1532 Vκ=3.5%, DH1533 V_H_=4.2%, DH1533 Vκ=1.5%).

**Figure 1:**
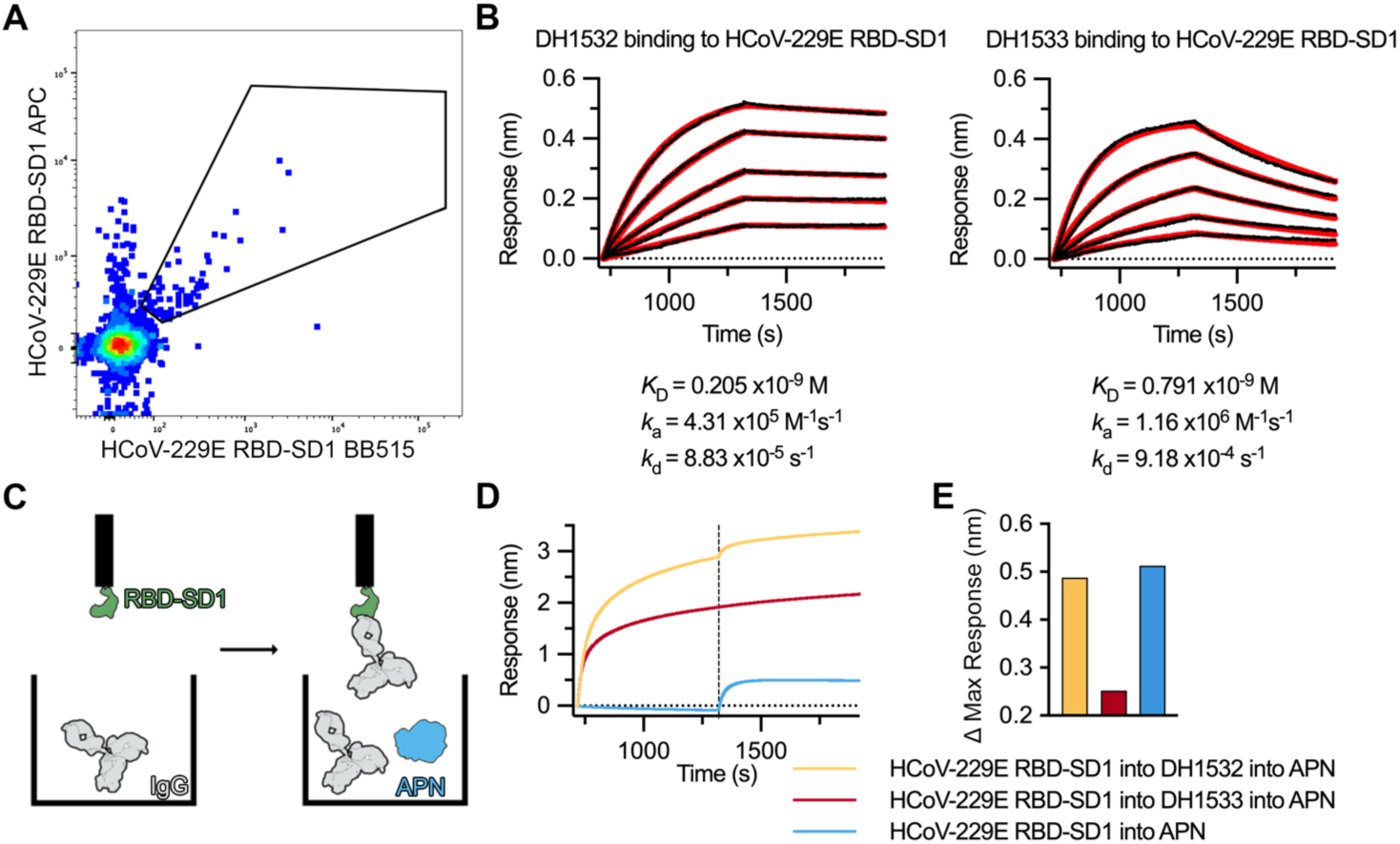
Isolation and characterization of HCoV-229E RBD-SD1-specific human antibodies. (**A**) A FACS dot plot of IgD^−^ B cells is shown, with cells stained using HCoV-229E RBD-SD1 conjugated to two different fluorochromes. (**B**) BLI sensorgrams measuring binding interactions for DH1532 (*left*) and DH1533 (*right*) to HCoV-229E RBD-SD1. Data are shown as black lines and 1:1 binding model fits are shown as red lines. The corresponding kinetic values for each interaction are shown below. (**C**) Cartoon depicting BLI-based receptor competition assay. Biosensors loaded with HCoV-229E RBD-SD1 were first dipped into wells containing IgG before being transferred to a well containing both IgG and soluble APN ectodomain. (**D**) Sensorgram showing results of receptor competition assay. The yellow biosensor was dipped into DH1532 during the first phase. The red biosensor was dipped into DH1533 during the first phase. The blue biosensor was dipped into buffer alone during the first phase. All biosensors were then transferred to wells containing APN during the second phase. (**E**) The difference in maximum response values between the first and second phases of the BLI experiment shown in **D** are plotted for each biosensor. The color of each bar is shared with the corresponding sensorgrams in **D**.

In the context of other coronaviruses, RBD-directed antibodies that disrupt receptor engagement oftentimes neutralize by either preventing viral interactions with susceptible host cells and/or by prematurely triggering S to the postfusion conformation. To determine whether these HCoV-229E RBD-directed mAbs might have the potential to neutralize by preventing the interaction between the RBD and APN, we performed a biolayer interferometry (BLI)-based receptor competition assay (**Figure 1C**). BLI biosensors were first loaded with HCoV-229E RBD-SD1 before being dipped into a well containing IgG for 600 seconds to allow antibody binding to occur. These biosensors were then dipped into another well containing both soluble APN and additional IgG to ensure that the RBD-SD1+IgG interaction was maintained throughout the course of the experiment. Upon exposure to APN, the biosensor containing RBD-SD1+DH1532 showed an additional increase in response, indicating that DH1532 binding did not prevent the association between the RBD and APN (**Figure 1D-E**). Under identical conditions, the DH1533-bound biosensor showed no such increase, suggesting that this antibody may neutralize HCoV-229E by directly disrupting receptor engagement.

Virus neutralization assays were next performed to more directly test the functional activity of these mAbs. Our group recently generated reverse genetic systems for the historical 1962 HCoV-229E VR-740 isolate and a contemporary clinical isolate from 2022, HCoV-229E UNC/2/2022, facilitating the creation of infectious nanoluciferase (nLUC) reporter viruses. The S proteins of VR-740 and UNC/2/2022 are 93.3% identical over 1155 residues, with 32 of these 79 differing amino acids residing in the RBD. Having generated nLUC reporter viruses for each, we then compared the neutralization capacity of DH1532 and DH1533 against recombinant HCoV-229E VR-740 and HCoV-229E UNC/2/2022 nLUC viruses in Caki1 cells. In agreement with our biophysical data, we found that DH1533 potently neutralized the contemporary HCoV-229E UNC/2/2022 nLUC virus with an IC_50_ of 0.62 μg/mL and as expected based on its failure to block APN binding, DH1532 exhibited weaker neutralization (IC_50_ = 8.67 μg/mL) (**Figure 2**). Efforts to determine the binding epitope of DH1532 by negative-stain electron microscopy (nsEM) ultimately yielded particles that resembled misfolded trimers rather than intact complexes of prefusion HCoV-229E S bound by Fab, indicating that DH1532 may recognize a cryptic epitope (**Supplementary Figure 2**). Neither mAb demonstrated the capacity to reach 50% neutralization of the historical VR-740 strain at the highest concentration tested (25 μg/mL). Thus, we demonstrate DH1533 neutralized contemporary but not antiquated HCoV-229E, confirming the importance of using more current, biologically relevant strains when evaluating immune responses and immunotherapies.

**Figure 2:**
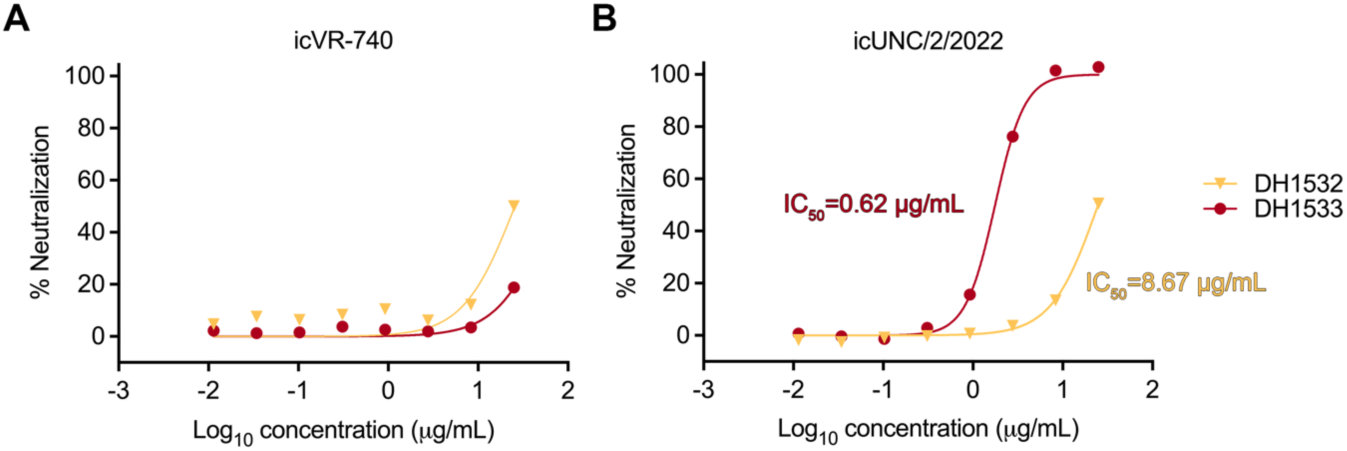
Strain-specific neutralization of HCoV-229E reporter viruses. Antibody neutralization activity against (**A**) icVR-740 and (**B**) icUNC/2/2022 reporter viruses. DH1532 is colored yellow and DH1533 is colored red. IC_50_ values for each antibody are labeled in the corresponding colors. Data are representative of two technical replicates.

Given the ability of DH1533 to potently neutralize the more contemporary HCoV-229E clinical isolate, but not the ancestral VR-740 strain, we performed a series of binding experiments using seven RBD-SD1 proteins (residues 278-497, VR-740 numbering), five of which were derived from the historically representative HCoV-229E strains proposed by Eguia et al^28^. The seven constructs, spanning viral evolution from 1984 to 2022, reflect overall shifts in spike antigenicity in the sense that the majority of mutations tend to accumulate in the receptor binding loops (residues 309-320, 355-358, 402-407) that mediate contact to APN (**Figure 3A, Supplementary Figure 3**). As has been reported previously, the sequence similarity between the RBDs of different variants directly correlates with the number of years between virus isolation^28,31^. For example, the RBD from the from the BN1/GER/2015 strain (2015) that was used for antigen-specific B cell isolation is 96.4% identical to the RBD from the Seattle/USA/SC677/2016 strain (2016), but only 79.9% identical to the RBD from the VR-740 strain (1962) (**Figure 3B**). DH1533 Fab exhibited robust binding to RBD-SD1 constructs from 2016 and 2015, but showed no detectable binding to RBD-SD1s from 1984-2008 at a concentration of 10 nM (**Figure 3C**). As expected based on our observed neutralization, DH1533 Fab also bound to the RBD-SD1 construct from 2022, albeit with slightly lower affinity (1.47 nM). This stark strain-restricted reactivity indicates that DH1533 recognizes an epitope that is subject to antigenic drift and suggests that it may have been elicited in response to more recent HCoV-229E infection, as opposed to prior exposure.

**Figure 3:**
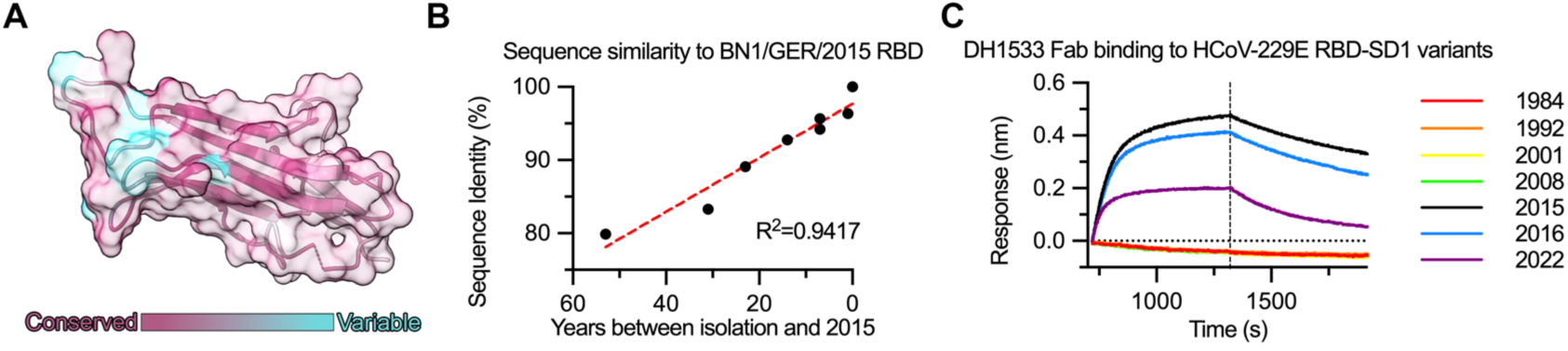
Viral sequence diversity in the RBD impacts DH1533 binding. (**A**) Sequence variability has been mapped onto the structure of the HCoV-229E RBD (PDB ID: 6U7G) using ConSurf. More conserved residues are colored purple and less conserved residues are colored teal. (**B**) Sequence identity of the RBD relative to the BN1/GER/2015 strain is plotted for representative HCoV-229E variants spanning 1962-2022. Time separating 2015 from the year when a strain was isolated is plotted on the x-axis. The red dashed line shows a linear regression, with an R^2^ value of 0.9417. (**C**) BLI sensorgrams are shown, measuring binding of DH1533 Fab to HCoV-229E RBD-SD1 constructs from representative strains isolated between 1984 and 2022.

To more precisely characterize the molecular basis for DH1533 binding and neutralization, we determined the cryo-EM structure of DH1533 Fab bound to a 2P-stabilized HCoV-229E S ectodomain spanning residues 1-1054 (HCoV-229E S2P). In contrast to most of the published cryo-EM structures of HCoV-229E spike^17,20^, we observed the S trimer in the compact “conformation 1”^32^, with the C-termini of all three protomers remaining tightly associated. Three copies of DH1533 Fab were observed binding to the S trimer perpendicular to the central axis, forming a C3-symmetrical, propellor-shaped complex (**Figure 4A, Supplementary Figures 4-5, Supplementary Table 1**). DH1533 binds to the solvent-accessible outer face of the beta sandwich that acts as a scaffold for the receptor binding loops at the tip of the RBD. Alignment of the DH1533-bound RBD to previously reported structures of the RBD bound to APN reveals that the DH1533 light chain framework would form large steric clashes with APN, providing a structural rationale for competition and antibody-mediated neutralization (**Figure 4B**). The precise contacts that mediate DH1533 binding are distributed somewhat evenly throughout the light chain and the heavy chain (**Figure 4C-D**), although the elongated 20 amino acid CDRH3 appears to play a critical role by wedging underneath the RBD such that Thr96, Arg99 and Tyr100b can form hydrogen bonds with the RBD residues Arg357, Asp351 and Ser361, respectively (**Figure 4D**). The light chain is positioned closer to the receptor binding loops at the tip of the RBD, and it forms extensive contacts with the positively charged RBD residues Arg311, Lys315 and Lys354, all of which lie within receptor binding loops 1 and 2.

**Figure 4:**
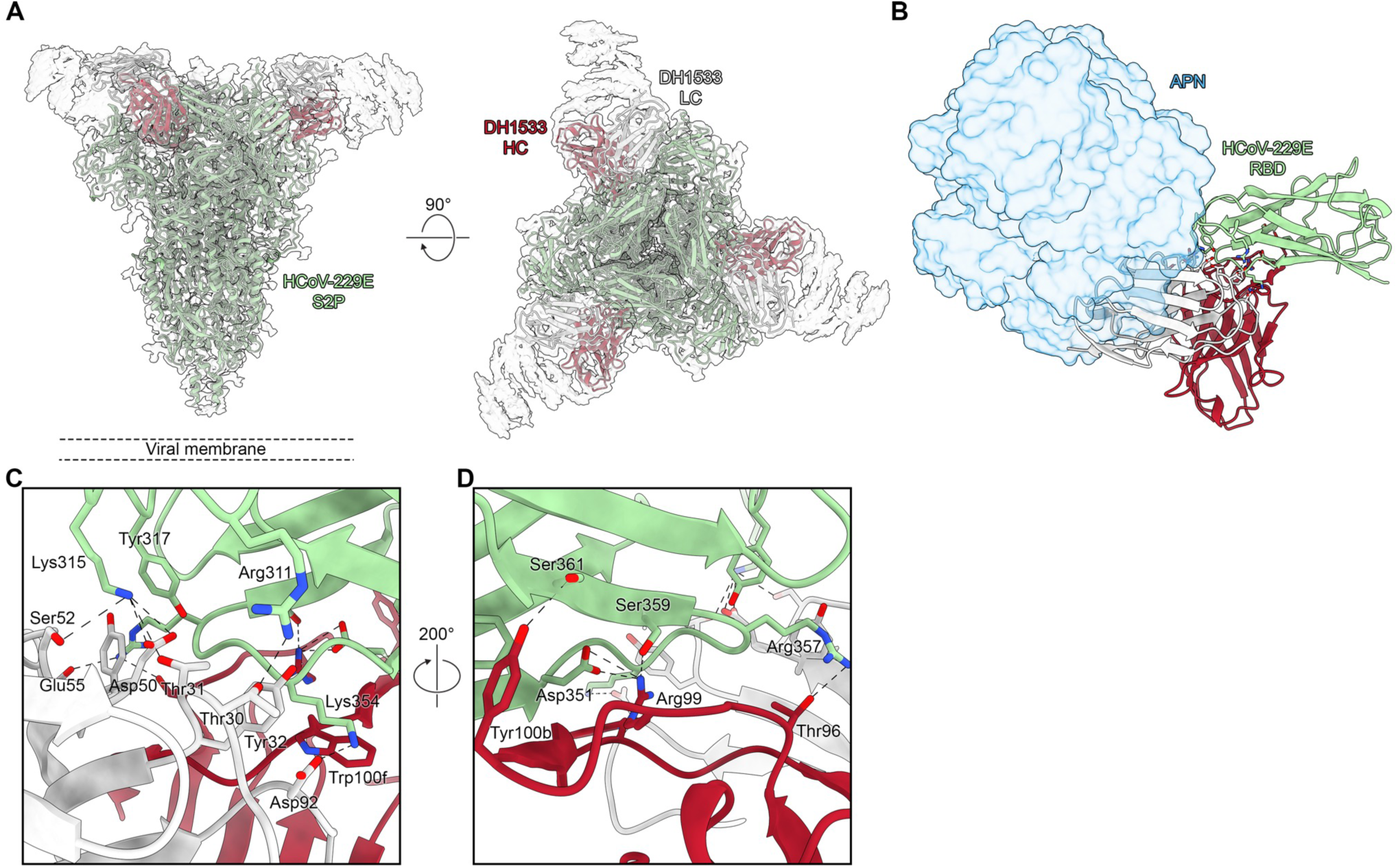
The cryoEM structure of DH1533 bound to HCoV-229E S. (**A**) The structure of DH1533-bound HCoV-229E S2P is shown as ribbon diagrams, docked into the corresponding transparent cryo-EM map from “side” (*left*) and “top” (*right*) views. HCoV-229E S2P is colored green, the DH1533 heavy chain is colored red and the DH1533 light chain is colored white. (**B**) An individual DH1533-bound RBD is shown in ribbon diagrams, with the APN ectodomain from a previously determined co-crystal structure (PDB ID: 6U7G), shown as a a blue, transparent molecular surface docked into the bound conformation relative to the HCoV-229E RBD. (**C**) A zoomed-in view of the DH1533 binding interface from the “top” view is shown. Chains are colored according to panel A and critical contact residues are shown as sticks. Oxygen atoms are colored red, nitrogen atoms are colored blue and predicted hydrogen bonds are shown as black dashed lines. (**D**) An additional view of the DH1533 binding interface is shown, rotated 200° relative to panel **C**.

These contact residues revealed by our cryo-EM structure also help to explain the strain-restricted binding and neutralization that were previously observed (**Supplementary Figure 6**). For example, the arginine at position 311 in the BN1/GER/2015 strain is a serine in the VR-740, 1984 and 1992 isolates. Similarly, these more ancestral viruses contain asparagine instead of lysine at position 345 and alanine or glycine residues instead of aspartic acid at position 351, all of which are predicted to eliminate key contacts between the RBD and DH1533. Other mutations, such as K315R and Y317F are more biochemically conservative, but may still play a role in reduced antibody recognition and antigenic drift, particularly when combined with other neighboring mutations within the same epitope.

In addition to the triply bound structure, two additional conformations could be resolved from the cryo-EM dataset. These reconstructions had either one or two copies of the DH1533 Fab bound and unexpectedly, the unbound protomers in these reconstructions displayed obvious conformational flexibility in the RBD (**Figure 5**). These unbound RBDs have rotated away from the NTD and S2 where they normally reside in the receptor inaccessible down conformation, undergoing a similar trajectory to the “up” and “down” kinetics that have been extensively characterized in the *Betacoronavirus* RBDs. The local resolution of these unbound RBDs is worse than 6 Å, even after extensive 3D variability analysis, particle subtraction and local refinement efforts, suggesting that this unmoored RBD is extremely conformationally flexible in the absence of a stabilizing binding partner such as APN. Similar to other previous cryo-EM structures of unbound HCoV-229E S, low-resolution cryo-EM analysis of HCoV-229E S2P in the absence of DH1533 Fab showed no evidence of RBD exposure (**Supplementary Figure 7**), suggesting that DH1533 binding is inducing this conformational flexibility in neighboring, unbound protomers. It should also be noted that we did not observe any unbound RBDs in the down conformation, likely reflecting the high binding affinity of DH1533 and the molar excess of Fab that was used to prepare the cryo-EM sample.

**Figure 5:**
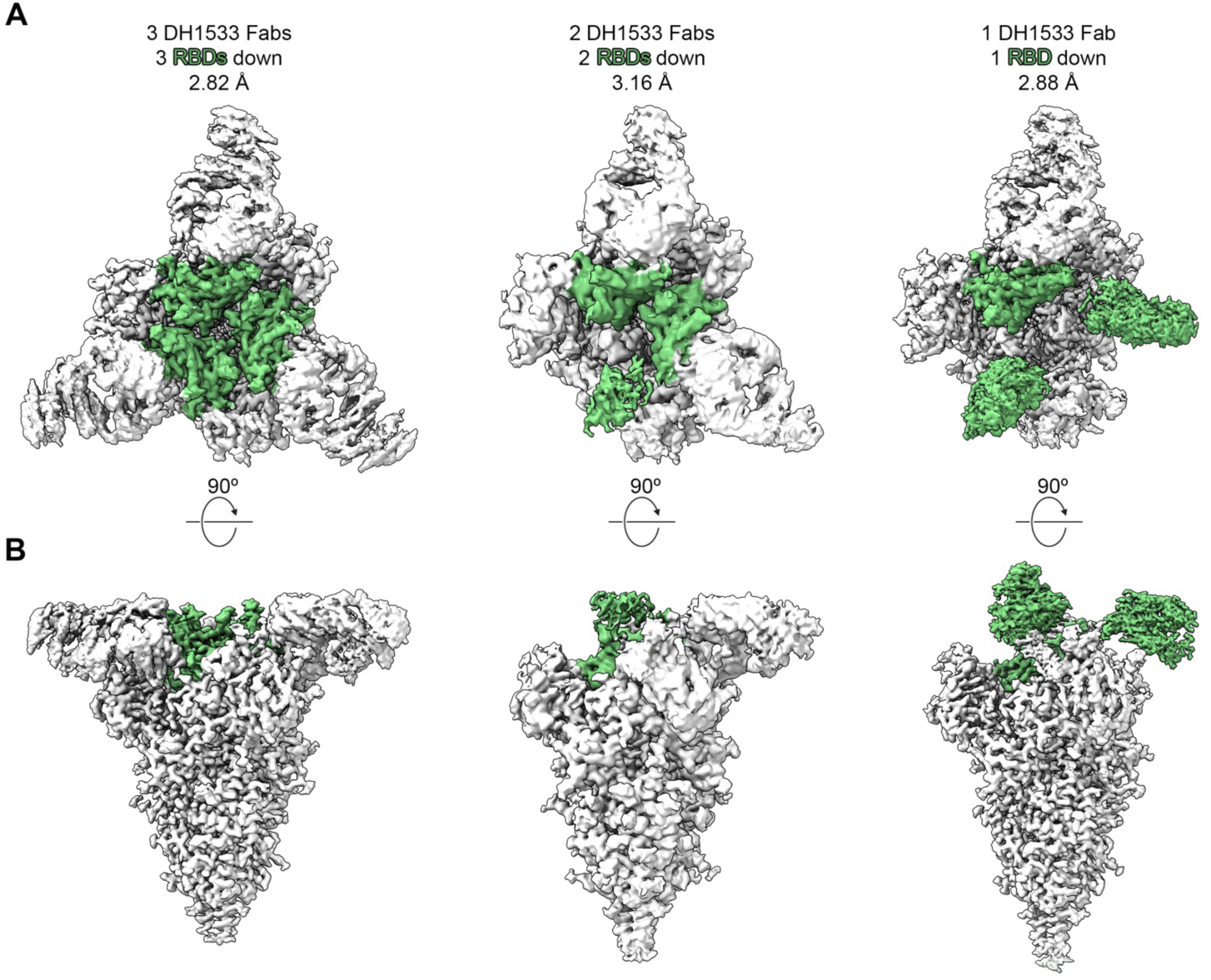
DH1533 binding induces RBD movement in neighboring, unbound protomers. (**A**) Top and (**B**) side views of the three cryo-EM reconstructions that were calculated from the DH1533 Fab + HCoV-229E S2P dataset are shown. The portions of each map corresponding to the RBDs have been colored green.

Comparison of the DH1533-bound protomer with previously reported cryo-EM structures of unbound HCoV-229E S protomers reveals that DH1533 binding causes the NTD of the bound protomer to shift downwards towards the viral membrane (**Supplementary Movie 1**). This downwards shift is likely a compensatory movement to minimize clashes between the DH1533 Fab and nearby *N*-linked glycans at positions 62, 98 and 176 within the NTD. It is possible that this downward shift of the NTD is then sufficient to allosterically release the neighboring RBDs from their receptor-inaccessible conformation, potentially leading to premature triggering of S even in the absence of host cell receptor. The observation of DH1533-bound dissociated S1 subunits within our cryo-EM dataset provides further support for this proposed mechanism (**Supplementary Figure 8**). Collectively, these data suggest that DH1533 is capable of neutralizing HCoV-229E via two overlapping methods. First is the relatively straightforward occlusion of the receptor binding loops, preventing the association of the host cell receptor APN. Additionally, DH1533 binding likely forces neighboring RBDs into the receptor accessible conformation, functionally acting as a receptor mimic to prematurely induce dissociation of the S1 subunit.

## DISCUSSION

Although HCoV-229E is thought to have first emerged into the human population approximately two hundred years ago^14,33^, little is known about how the humoral immune system responds to this pervasive pathogen. By probing PBMC samples from a donor convalescing after HCoV-229E infection, in these studies we have isolated the first human HCoV-229E S-directed antibodies. Both of these antibodies demonstrated the capacity to neutralize a contemporary HCoV-229E clinical isolate, although DH1533 showed greater potency than DH1532. Curiously, extended incubation of DH1532 Fab with the soluble HCoV-229E S2P ectodomain yielded dissociated protein particles by nsEM analysis, reminiscent of the effect that the mAb CR3022 has on the *Betacoronavirus* SARS-CoV-2 spike^34^. CR3022 is known to bind a cryptic epitope on the inner face of the SARS-CoV-2 RBD that is not accessible in the prefusion conformation of S. Likewise, the HCoV-229E RBD-directed mAb S11, which was generated by panning of a synthetic phage display library, binds to an epitope near the base of the RBD which is not accessible in the context of the prefusion HCoV-229E spike^35^. Nevertheless, S11 has been shown to neutralize HCoV-229E pseudovirus with an IC_50_ value of 19 μg/mL, suggesting that it may function by destabilizing the prefusion trimer and rendering it non-fusogenic. Given our nsEM and viral neutralization data, it seems likely that DH1532 functions in a similar way, perhaps after being elicited by dissociated S1 that is shed during natural infection^36^.

DH1533 neutralizes with far greater potency and does so by both blocking the interaction with APN and by functioning as a receptor mimic^37^ (**Figures 1D, 2, 4B**). However, despite its potent neutralization of our HCoV-229E 2022 clinical isolate, it fails to neutralize the ancestral VR-740 strain, reflecting the antigenic drift of HCoV-229E over time which has been investigated using historical sera samples and pseudoviruses spanning HCoV-229E strains from 1984-2016^28^. By testing binding to HCoV-229E RBD-SD1 constructs ranging from 1984-2022, we show that DH1533 fails to bind to RBDs from strains prior to 2015, suggesting that although HCoV-229E may evolve at slower rates than influenza^38^ or *Betacoronaviruses* like HCoV-OC43^39^ or SARS-CoV-2^40^, it is still ultimately capable of evading neutralizing antibodies through continuous antigenic drift (**Figure 3C**). We observed higher affinity DH1533 Fab binding to RBD-SD1 constructs from 2015 and 2016 viruses, compared to the RBD-SD1 from the more contemporary 2022 clinical isolate. These data suggest that DH1533 may have initially been elicited in response to prior HCoV-229E infection, although the lack of longitudinal sampling makes it difficult to investigate this phenomenon more definitively. However, by using 2015 RBD-SD1 tetramers to isolate antigen-specific B cells, we may have selectively enriched for B cells that were reactivated in response to this more recent HCoV-229E infection. These findings have critical implications for the design of future vaccine immunogens, suggesting that in order to elicit protective immune responses, immunogens must either focus the immune response towards more conserved epitopes like the S2 subunit^41^, or they may need to be iteratively updated to reflect gradual changes in the antigenicity of circulating viral strains.

The cryo-EM structure of HCoV-229E S2P bound by three DH1533 Fabs revealed a C3-symmetrical complex, with DH1533 binding to the outer face of the RBD in the down conformation. The position of the DH1533 light chain near the distal receptor binding loops of the RBD causes steric hindrance that prevents APN association^22^, which we propose is the major mechanism of HCoV-229E neutralization. Analysis of the non-symmetrical cryo-EM structures of DH1533 bound to HCoV-229E S2P also revealed that DH1533 binding is capable of allosterically inducing movement of neighboring, unbound RBDs out of the receptor-inaccessible down conformation (**Figure 5**). In some ways, this is similar to the recently reported structure of the HCoV-229E S ectodomain bound to an APN homodimer^20^. However, whereas the APN-bound up RBD could be resolved relatively clearly, the movement that we observe appears to be much less coordinated, and the portions of our cryo-EM maps corresponding to the up RBDs quickly disappear at higher thresholds, preventing us from building models of this transient conformational intermediate. Whether this extreme conformational flexibility is an inherent property of the up RBD when it is not stabilized by APN binding or whether it is a non-physiological conformation that is the result of nearby DH1533 binding remains unclear and requires additional investigation.

The emergence potential of the CoV family is undeniable given that at least three novel HCoVs have emerged in the past two decades. Furthermore, this phenomenon is not unique to the *Betacoronaviruses*, as evidenced by recent sporadic human infections caused by zoonotic *Alphacoronaviruses*, including zoonotic strains that replicate in human primary cells from the lungs and intestine^11,12^. Thus, the study of *Alphacoronavirus* immunity and the development of therapeutics is essential to prepare for potential future novel *Alphacoronavirus* emergence. Overall, these studies demonstrate that the humoral immune system can produce high-affinity HCoV-229E RBD-directed antibodies that exhibit potent neutralization even after relatively minimal somatic hypermutation. These findings indicate that the creation of effective prophylactics and therapeutics to reduce HCoV-229E disease burden are possible, although the threat of gradual viral escape via antigenic drift necessitates careful consideration of evolving global sequence variation. Nevertheless, we provide important insights into CoV type-specific immunity which could inform next generation, broadly acting CoV medical countermeasures.

## AUTHOR CONTRIBUTIONS

E.B.W., K.O.S., R.S.B., B.F.H., D.W.C., T.P.S. and D.W. designed the research. M.F.L., T.S., H.M.F., C.P., B.L.Y., Q.Y., T.N.S., R.J.E. and D.W. performed research. R.J.E., D.W.C., T.P.S. and D.W. analyzed data. T.P.S. and D.W. wrote the manuscript with input from all authors.

## ACKNOWLEDGEMENTS

The APN ectodomain expression plasmid was a generous gift from Dr. Jason S. McLellan. Cryo-EM studies were performed at the Duke University Shared Materials Instrumentation Facility (SMIF), a member of the North Carolina Research Triangle Nanotechnology Network (RTNN), which is supported by the National Science Foundation (ECCS-2025064) as part of the National Nanotechnology Coordinated Infrastructure (NNCI). Flow cytometry was performed at the DHVI Flow Cytometry Facility, which is supported in part by the National Institute of Allergy and Infectious Diseases (UC6-AI058607). This project was supported by the NIAID award P01 AI158571 (to B.F.H.).

## MATERIALS AND METHODS

### Human donor sample collection

Human subject studies were approved by the Duke University Health System Institutional Review Board (IRB) and conducted in agreement with the policies and protocols approved by the Duke IRB, consistent with the Declaration of Helsinki. Written informed consent was obtained from the research subject or their legally authorized representatives. A nasopharyngeal swab was collected and analyzed using the QIAstat-DX Respiratory Panel Plus cartridge within the QIAstat-DX Analyzer 1.0 RT-PCR system (QIAGEN). PBMC samples were collected from the same participant 2 weeks after the onset of symptoms.

### Protein expression and purification

Plasmids encoding residues 1-1054 of HCoV-229E S (BN1/GER/2015) with 2P substitutions^27^ at positions 869 and 870, a C-terminal T4 fibritin motif, an HRV3C cleavage site, an 8× HisTag and a TwinStrepTag; human aminopeptidase N residues 66-967 with an artificial signal peptide, a C-terminal HRV3C cleavage site, a monomeric human Fc tag and an 8× HisTag; HCoV-229E RBD-SD1 residues 277-494 (VR-740 numbering) from strains HCoV-229E-5/9/84, HCoV-229E-17/6/92, HCoV-229E-8/8/01, 693A_2008, Seattle/USA/SC677/2016, BN1/GER/2015, UNC/2/2022 and VR-740 with an artificial signal peptide, a C-terminal HRV3C cleavage site, a monomeric human Fc tag and an 8× HisTag; HCoV-229E RBD-SD1 residues 277-495 (BN1/GER/2015) with an artificial signal peptide, a C-terminal HRV3C cleavage site, an 8× HisTag and an Avi tag were transfected into FreeStyle293 cells using polyethylinimine. Plasmids encoding for furin and for DS-Cav1 with a C-terminal T4 fibritin motif, an HRV3C cleavage site, an 8× HisTag and an AviTag were cotransfected using the same conditions. Similarly, plasmids encoding for the heavy and light chains of DH1532 and DH1533 IgG with an HRV3C protease cleavage site engineered into the hinge between the CH1 and CH2 domains of the heavy chain were cotransfected using the same conditions.

HCoV-229E S2P was purified from cell supernatants using StrepTactin resin (IBA) before being run over a Superose 6 Increase 10/300 GL column in buffer composed of 2 mM Tris pH 8.0, 200 mM NaCl and 0.02% NaN_3_. HCoV-229E RBD-SD1 monoFc constructs, APN monoFc, DH1532 IgG and DH1533 IgG were purified from cell supernatants using Protein A resin before being run over a Superdex 200 Increase 10/300 GL column in buffer composed of 2 mM Tris pH 8.0, 200 mM NaCl and 0.02% NaN_3_. HCoV-229E RBD-SD1 HisAvi and DS-Cav1 HisAvi were purified from cell supernatants using NiNTA resin before being run over a Superdex 200 Increase 10/300 GL column in PBS. APN monoFc, DH1532 IgG and DH1533 IgG were treated with HRV3C protease overnight at 4 °C to cleave C-terminal affinity tags and to separate Fabs from Fc domains. Untagged APN was then purified by SEC using a Superdex 200 Increase 10/300 GL column in buffer composed of 2 mM Tris pH 8.0, 200 mM NaCl and 0.02% NaN_3_. DH1533 and DH1532 Fabs were first purified using CaptureSelect IgG-CH1 affinity resin (ThermoScientific) to remove excess protease and cleaved Fcs before a final SEC purification using a Superdex 200 Increase 10/300 GL column in buffer composed of 2 mM Tris pH 8.0, 200 mM NaCl and 0.02% NaN_3_.

### Antigen-specific B cell sorting

HCoV-229E RBD-SD1 HisAvi and DS-Cav1 HisAvi were biotinylated by BirA treatment for 4 hours at 30 °C. Biotinylated proteins were then purified by SEC using a Superdex 200 Increase 10/300 GL to remove excess free biotin. Biotinylated HCoV-229E RBD-SD1 was then tetramerized by the addition of either streptavidin-BB515 or streptavidin-APC. Biotinylated DS-Cav1 was tetramerized by the addition of streptavidin-BV421.

Human donor PBMCs were stained with PE anti-human IgD (clone IA6-2, BD Biosciences), PE-Cy5 anti-human CD235a (clone GA-R2, BD Biosciences), PE-Cy5 anti-human CD3 (clone HIT3a, BD Biosciences), (PE-Cy7 anti-human CD27 (clone O323, ThermoFisher), APC A700 anti-human CD38 (clone LS198-4-3, Beckman Coulter), APC-Cy7 anti-human CD19 (clone SJ25C1, BD Biosciences), BV570 anti-human CD16 (clone 3G8, Biolegend), BV605 anti-human CD14 (clone M5E2, Biolegend), BV711 anti-human IgM (clone G20-127, BD Biosciences), and fluorescently-labeled viral glycoproteins (HCoV-229E RBD-SD1-BB515, HCoV-229E RBD-SD1-APC, and DS-Cav1-BV421). Cells were further labeled with Fixable Aqua Live/Dead Cell Stain Kit (Invitrogen). Antigen-specific (HCoV-229E RBD-SD1^+/+^/DS-Cav1^−^) memory B cells were identified as viable CD3^−^/CD14^−^/CD16^−^/CD235a^−^/CD19^+^/IgD^−^/probe^+^ cells on a BD FACSymphony S6 flow cytometer (BD Biosciences) and single cells were sorted into a 96-well plate containing lysis buffer. The full cell gating strategy is depicted in **Supplementary Figure 1**. The collection plate containing individual sorted cells was then rapidly frozen in a dry ice and ethanol slurry before being stored at −80°C. Flow cytometric data were analyzed using FlowJo v10.10 (BD Biosciences).

### Immunoglobulin gene cloning by PCR and preliminary mAb screening

Immunoglobulin genes were amplified from antigen-specific, sorted single B cells by RT-PCR using methods described previously^42,43^, with minor modifications. Briefly, immunoglobulin genes from single B cells were reverse transcribed using Superscript III (ThermoFisher) and random hexamer primers (GeneLink). The resulting cDNA was then used as a template for two rounds of nested PCR using both constant and variable region primers. PCR products were then sequenced using both forward and reverse primers that recognized tags introduced during the second round of PCR. Forward and reverse immunoglobulin gene sequences were assembled using the Duke Human Vaccine Institute Automated Sequence Analysis Pipeline (DHVI ASAP). Sequences were subjected to IGHV, IGKV and IGLV sequence assignment and mutation frequency analysis using the human library in Cloanalyst^44^. Productive antibody gene pairs were synthesized and cloned into gamma, kappa, or lambda expression plasmids that were then co-transfected into Expi293 cells using polyethylinimine. After five days, cell supernatants were harvested and tested for binding reactivity to HCoV-229E RBD-SD1 HisAvi by ELISA. HCoV-229E RBD-SD1 HisAvi was directly coated to a 384 well plate at 2 μg/mL overnight before being blocked with PBS containing 4% whey protein, 15% goat serum, 0.5% Tween 20 and 0.05% NaN_3_. Supernatants were then incubated for 90 minutes before the plate was washed and binding was detected with goat anti-human Fc HRP (Jackson ImmunoResearch) and TMB substrate (Sera Care Life Sciences).

### Binding kinetics determination by biolayer interferometry

DH1532 IgG and DH1533 IgG were immobilized to anti-human capture (AHC) biosensors (Sartorius) to a response level of 1.6 nm over the course of 600 seconds. Biosensors were then dipped into wells containing two-fold dilution series of HCoV-229E RBD-SD1 HisAvi ranging from either 10−0.625 nM (DH1532) or 5−0.3125 nM (DH1533) for 600 seconds before being transferred to wells containing empty buffer for 600 seconds. Data were reference-subtracted and processed using Octet Data Analysis software v12.0 (Sartorius) with a 1:1 binding model.

### Biolayer interferometry receptor-competition assay

HCoV-229E RBD-SD1 HisAvi (BN1/GER/2015) was immobilized to NiNTA biosensors (Sartorius) to a response level of 1.5 nm over the course of 600 seconds. Biosensors were then dipped into wells containing either 100 nM DH1532 IgG, 100 nM DH1533 IgG or running buffer alone for 600 seconds. Sensors were then dipped into wells containing either 100 nM DH1532 IgG + 100 nM APN, 100 nM DH1533 IgG + 100 nM APN or 100 nM APN alone for 600 seconds. Data were reference-subtracted and processed using Octet Data Analysis software v12.0 (Sartorius).

### Reporter virus production and neutralization

icHCoV 229E VR740 and icHCoV 229E UNC/2/2022 viruses were engineered to express nanoluciferase (nLUC) in place of ORF 4 and were recovered via reverse genetics using general methods previously described^45^. Infectious viral titers were measured in CaKi-1 cells via plaque assay in 6 well plate format and in triplicate biological replicates and expressed as plaque forming units per mL (PFU/mL). For the neutralization assay, CaKi-1 cells were plated at 3 × 10^4^ cells per well in black bottomed, black walled plates the day prior to the assay. mAbs DH1532 and DH1533 were diluted in CaKi-1 infection media (Corning McCoy’s 5A Iwakata & Grace modified media supplemented with 5% FBS, 1x Gibco GlutaMax, 1x Gibco NEAA, and 1x Gibco Anti/Anti) starting at a concentration of 50 µg/mL and serially diluted in 3-fold steps 7 times. Serially diluted mAbs were mixed in equal volume with diluted virus (1.6 × 10^4^ PFU/mL). Antibody-virus and virus only mixtures were incubated at 32°C with 5% CO_2_ for 1 hour and following incubation, added in duplicate to cells at a final concentration range of 25 to 0.01 µg/mL of mAb and 800 PFU/well of virus. The same concentration of virus was added to cell-free wells to control for background caused by virus stock associated luciferase. Cells were incubated at 32°C with 5% CO_2_ and after 43 hours, viral replication was measured via Nano-Glo Luciferase Assay System (Promega, Cat.#: N1150) on a GloMax Explorer Multimode Microplate Reader (Promega, Madison, WI). Percent neutralization was calculated by determining the percent reduction of RLUs relative to the average of virus-only infected control wells. Neutralization curves were generated using GraphPad Prism software version 10.4.2. The details of the generation and characterization of the wildtype and derivative nLUC and GFP reporter, time-ordered HCoV-229E molecular clones will be reported in future publications (Liccione and Pajon et al., *manuscript in preparation*).

### Negative stain EM

A frozen aliquot of HCoV-229E S2P was thawed for 20 mins at 37 °C and DH1532 Fab was thawed at 4 °C. DH1532 Fab was mixed with spike at a molar ratio of 15:1 Fab to spike. The mixture was incubated overnight at 4 °C to allow binding to occur. Sample was then diluted to 400 µg/mL in 20 mM HEPES, 150 mM Na_2_SO_4_, 2.5 mM NaCl, 5 mM NaN_3_ pH 7.4 (HBSO_4_-150) buffer containing 8 mM glutaraldehyde. After a 5 minute incubation, glutaraldehyde was quenched by the addition of 1 M Tris, pH 7.4, for a final concentration of 80 mM Tris. The quenched sample was then incubated at room temperature for an additional 5 minutes. Quenched sample was then diluted with HBSO_4_-150 for a final concentration of 50 µg/mL. Sample was then applied to a glow-discharged carbon-coated EM grid for 10-12 seconds, before excess liquid was removed by blotting, and 2 g/dL uranyl formate stain was applied for 1 minute. Stain was removed by blotting and the grid was air-dried. The stained grid was inspected on a Philips EM420 microscope operating at 120 kV and a nominal magnification of 49,000x. A total of 45 images were collected on a 76 Mpix CCD camera at 2.4 Å/pix. Images were processed and analyzed by 2D classification in Relion 3.0^46^.

### DH1533 binding to historical HCoV-229E RBD-SD1 variants

HCoV-229E RBD-SD1 monoFc variants were immobilized to anti-human capture (AHC) biosensors (Sartorius) to a response level of 0.9 nm over the course of 600 seconds. Biosensors were then dipped into wells containing 10 nM DH1533 Fab for 600 seconds before being transferred to wells containing empty buffer for 600 seconds. Data were reference-subtracted and processed using Octet Data Analysis software v12.0 (Sartorius).

### DH1533-bound HCoV-229E S2P cryo-EM data collection, processing and refinement

0.35 mg of HCoV-229E S2P was mixed with 0.13 mg of DH1533 Fab and binding was allowed to occur at room temperature for 20 minutes before 3 μL of protein mixture was applied to a glow-discharged Quantifoil Cu-200 1.2/1.3 grid. Excess liquid was blotted away for 5.5 seconds using a Vitrobot Mark IV (FEI) maintained at 4° C and 100% humidity. Data were collected using a Titan Krios G3i (ThermoFisher) operating at 300 kV with a K3 Bioquantum detector (Gatan) using Latitude S automation software (Gatan). 19,477 movies were collected at a calibrated magnification of 1.08 Å/pix. A full description of data collection parameters can be found in **Supplementary Table 1**.

Unaligned movies were imported into cryoSPARC v4.6.2 (Structura Biotechnology)^47^ for pre-processing with patch-based motion correction and patch-based CTF estimation. Blob-based particle picking was performed on a subset of movies to generate templates for particle picking by 2D classification. These 2D templates were then used to pick a total of 6,573,334 particles. 2D classification was performed to remove junk particles and the remaining particles (2,245,907) were used as inputs for iterative rounds of *ab initio* 3D reconstruction and 3D classification. Final particle stacks for each conformation were used as inputs for non-uniform 3D refinement^48^ and reconstructions were sharpened using DeepEMhancer^49^. Additional details regarding data processing are depicted in **Supplementary Figures 4 and 5**.

PDB ID: 7CYC^32^ was used as an initial model for HCoV-229E S2P and a model for the VH and VL domains of DH1533 was generated using AlphaFold2. Initial models were docked into 3D reconstructions, modified and iteratively refined using UCSF ChimeraX^50^, Coot^51^, ISOLDE^52^ and Phenix^53^. A full list of model statistics can be found in **Supplementary Table 1**.

### Unbound HCoV-229E S2P cryo-EM data collection and processing

3 μL of 0.35 mg of HCoV-229E S2P was applied to a glow-discharged Quantifoil Cu-200 1.2/1.3 grid. Excess liquid was blotted away for 6.0 seconds using a Vitrobot Mark IV (FEI) maintained at 4° C and 100% humidity. Data were collected using a Tundra Cryo TEM (ThermoFisher) operating at 100 kV with a Ceta-F CMOS detector using Smart EPU software (ThermoFisher). 191 micrographs were collected at a calibrated magnification of 1.21 Å/pix.

Micrographs were imported into cryoSPARC v4.6.2 (Structura Biotechnology)^47^ for CTF estimation using CTFFIND4^54^. Blob-based particle picking was performed to pick a total of 135,442 particles. After 2D classification, the remaining 23,808 particles were used as inputs for *ab initio* 3D reconstruction, 3D classification and the calculation of an 8.1 Å 3D reconstruction using 17,995 particles.

## DATA AVAILABILITY

All DH1533-bound cryo-EM reconstructions have been deposited to the Electron Microscopy Data Bank under accession codes EMD-70440, EMD-70441, EMD-70442, EMD-70507 and EMD-70508. The corresponding structural coordinates have been deposited to the Protein Data Bank under accession codes 9OFO, 9OFP and 9OFQ. All flow cytometry data are available upon request.

## FIGURE LEGENDS

**Supplementary Figure 1: Antigen-specific human B cell gating strategy**. (**A-I**) Each subsequent FACS plot displays only the population of cells that were included in the gate of the previous plot.

**Supplementary Figure 2: DH1532 destabilizes the HCoV-229E S2P trimer**. (**A**) A representative negative-stain electron micrograph of the HCoV-229E S2P trimer after overnight incubation with a molar excess of DH1532 Fab. (**B**) Corresponding 2D class averages of particles collected from the DH1532 Fab + HCoV-229 S2P mixture.

**Supplementary Figure 3: Sequence alignment of the HCoV-229E RBD**. Alignment of HCoV-229E RBD sequences from representative strains generated using Clustal Omega. Receptor binding loop residues have been highlighted blue.

**Supplementary Figure 4: Cryo-EM data processing workflow.**

**Supplementary Figure 5: Cryo-EM validation**. The FSC curves (*left*), viewing direction distribution plots (*center*) and local resolution maps (*right*) are shown for (**A**) triply bound HCoV-229E S2P, (**B**) doubly bound HCoV-229E S2P, (**C**) singly bound HCoV-229E S2P, and (**D**) a local refinement of the singly bound complex, focused on an “up” RBD and the nearby DH1533-bound RBD. (**E**, *left*) The molecular model of HCoV-229E S2P bound by a single DH1533 Fab is shown docked into the composite map generated from the reconstructions shown in **C** and **D**. Zoomed-in views of the same model and map are shown, highlighting the DH1533 heavy chain (*left*) and light chain (*right*).

**Supplementary Figure 6: Structural analysis of antigenic drift in the DH1533 epitope**. (A) The structure of the variable domain of DH1533 bound to the RBD is shown from two different angles in ribbon diagrams, colored according to **Figure 4**. Residues that vary between the representative strains in **Supplementary Figure 3** are colored purple. (B) A zoomed-in view of the DH1533 binding interface. Critical contact residues are shown as sticks. Oxygen atoms are colored red, nitrogen atoms are colored blue and predicted hydrogen bonds are shown as black dashed lines. Critical contact residue mutations have been labeled.

**Supplementary Figure 7: HCoV-229E S2P displays no conformational flexibility in the RBD in the absence of DH1533**. (**A**) Cryo-electron micrograph showing unbound HCoV-229E S2P trimers. (**B**) 2D class averages of HCoV-229E S2P particles. (**C**) An 8.1 Å 3D reconstruction of HCoV-229E S2P is shown from “side” and “top” views.

**Supplementary Figure 8: Dissociated S1 subunits observed by cryo-EM**. (**A**) 2D class averages from the DH1533 Fab + HCoV-229E S2P cryo-EM dataset that show dissociated S1 “rings”. Rings bound by a single DH1533 Fab have been highlighted yellow, rings bound by two DH1533 Fabs have been highlighted blue and rings with faint features suggesting three bound DH1533 Fabs have been highlighted purple. (**B**) Models corresponding to the 2D class averages in panel **A** are depicted, highlighted with the same coloring scheme. S1 ring models are colored green and DH1533 Fab models are colored red. (**C**) A 2D class average showing a “top” view of an intact, triply bound DH1533 Fab + HCoV-229E S2P complex has been highlighted green, with the corresponding model shown below in panel (**D**).

**Supplementary Movie 1: DH1533 binding causes compensatory movement in the neighboring NTD.** A single HCoV-229E S protomer is shown, interpolating the previously reported unbound conformation (PDB ID: 7CYC) and the DH1533-bound conformation. DH1533 binding causes the neighboring NTD to move downward, away from the RBD. The NTD is colored blue, the RBD is colored green and the rest of the protomer is colored white.

**Supplementary Table 1: Cryo-EM data collection and refinement statistics.**

