## Supplementary Information for "A neutralizing human antibody induces movement of the HCoV-229E receptor binding domain"

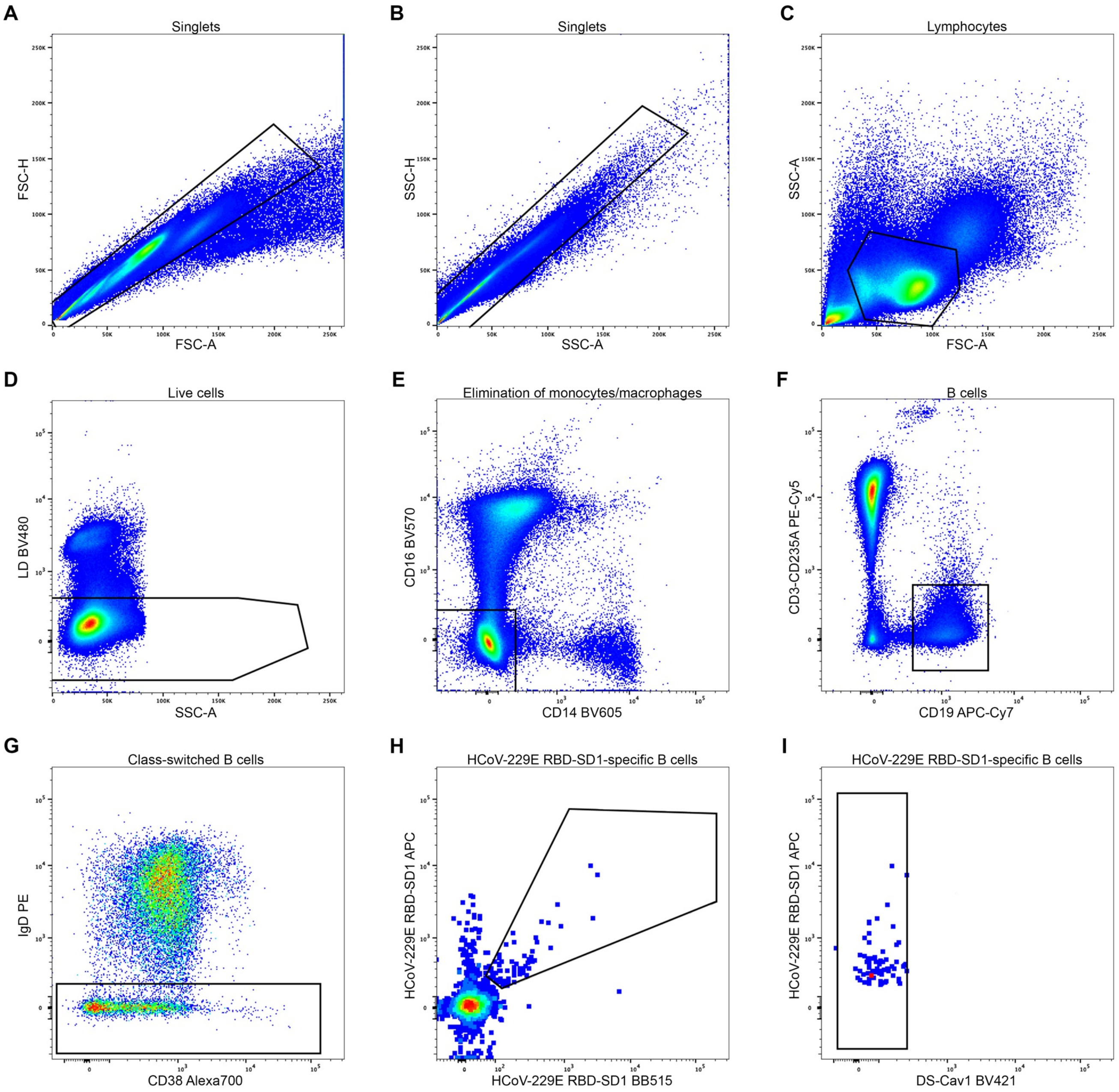


**Supplementary Figure 1: Antigen-specific human B cell gating strategy**. (**A-I**) Each subsequent FACS plot displays only the population of cells that were included in the gate of the previous plot.

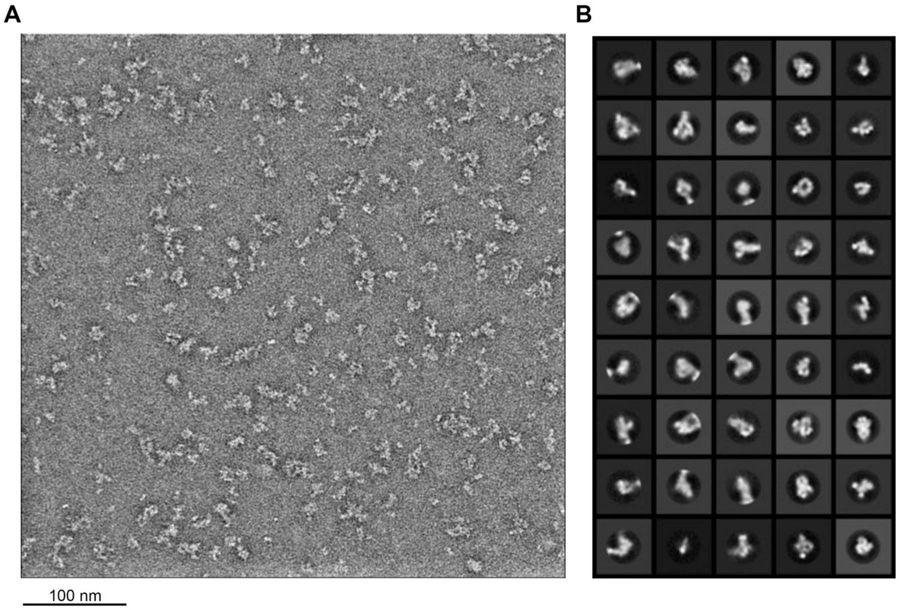


**Supplementary Figure 2: DH1532 destabilizes the HCoV-229E S2P trimer**. (**A**) A representative negative-stain electron micrograph of the HCoV-229E S2P trimer after overnight incubation with a molar excess of DH1532 Fab. (**B**) Corresponding 2D class averages of particles collected from the DH1532 Fab + HCoV-229 S2P mixture.

```
VR-740                       297 HKHTFIVLYVDFKPQSGGGKCFNCYPAGVNITLANFNETKGPLCVDTSHFTTKYVAVYAN 356
icUNC/2/2022                 296 HKHTFIVLHVKFEHGRGPGKCYNCRPAVINITLANFNETKGPLCVDTSHFTTQFVDN-VK 354
BN1/GER/2015                 296 HKHTFIVLYVNFEHRRGPGKCYNCRPAVINITLANFNETKGPLCVDTSHFTTQFVDN-VK 354
Seattle/USA/SSC677/2016      296 HKHTFIVLHVKFEHQRGPGKCYNCRPSVINITLANFNETKGPLCVDTSHFTTQFVDN-VK 354
HCoV-229E-5/9/84 (1984)      296 HKHTFIVLYVDFKLQSGVGRCFNCRPAVVNITLANFNETKGPLCVDTSHFTTKFVG--AN 353
693A_2008                    296 HKHTFIVLYVNFELRRGPGRCYNCRPAVINITLANFNETKGPLCVDTSHFTTQFVG--VK 353
HCoV-229E-17/6/92 (1992)     296 HKHTFIVLYVNFKLRSGVGRCYNCRPAVVNITLANFNETKGPLCVDTSHFTTQFVG--AK 353
HCoV-229E-8/8/01 (2001)      296 HKHTFIVLYVNFELRRGPGRCYNCRPAVVNITLANFNETKGPLCVDTSHFTTQFVG--VK 353
                                 ********:*.*:   * *:*:** *: :***********************::*   .:

VR-740                       357 VGRWSASINTGNCPFSFGKVNNFVKFGSVCFSLKDIPGGCAMPIVANWAYSKYYTIGSLY 416
icUNC/2/2022                 355 LARWSASITTGNCPFSFGKVNNFVKFGSVCFSLKAIPGGCAMPIMANLVNYKSHNIGSLY 414
BN1/GER/2015                 355 LARWSASINTGNCPFSFGKVNNFVKFGSVCFSLKDIPGGCAMPIMANLVNSKSHNIGSLY 414
Seattle/USA/SSC677/2016      355 LARWSASINTGNCPFSFGKVNNFVKFGSVCFSLKDIPGGCAMPIMANLVNHKSHNIGSLY 414
HCoV-229E-5/9/84 (1984)      354 FGRWSASINTGNCPFSFGKVNNFVKFGSVCFSLKDIPGGCAMPIVANLAYLNSYTIGSLY 413
693A_2008                    354 FDRWSASINTGNCPFSFGKVNNFVKFGSVCFSLKDIPGGCAMPIMANLVNHKSHNIGSLY 413
HCoV-229E-17/6/92 (1992)     354 FDRWSASINTGNCPFSFGKVNNFVKFGSVCFSLKDIPGGCAMPIMANLANLNSHTIGSLY 413
HCoV-229E-8/8/01 (2001)      354 FDRWSASINTGNCPFSFGKVNNFVKFGSVCFSLKDIPGGCAMPIMANLANLNSHNIGSLY 413
                                 . ******.************************* *********:** .  : :.*****

VR-740                       417 VSWSDGDGITGVPQPVEGV 435
icUNC/2/2022                 415 VSWSDGDVITGVPKPVEGV 433
BN1/GER/2015                 415 VSWSDGDVITGVPKPVEGV 433
Seattle/USA/SSC677/2016      415 VSWSDGDVITGVPKPVEGV 433
HCoV-229E-5/9/84 (1984)      414 VSWSDGDVITGVPKPVEGV 432
693A_2008                    414 VSWSDGDVITGVPQPVEGV 432
HCoV-229E-17/6/92 (1992)     414 VSWSDGDVITGVPKPVEGV 432
HCoV-229E-8/8/01 (2001)      414 VSWSDGDVITGVPKPVEGV 432
                                 ******* *****:*****
```

**Supplementary Figure 3: Sequence alignment of the HCoV-229E RBD**. Alignment of HCoV-229E RBD sequences from representative strains generated using Clustal Omega. Receptor binding loop residues have been highlighted blue.

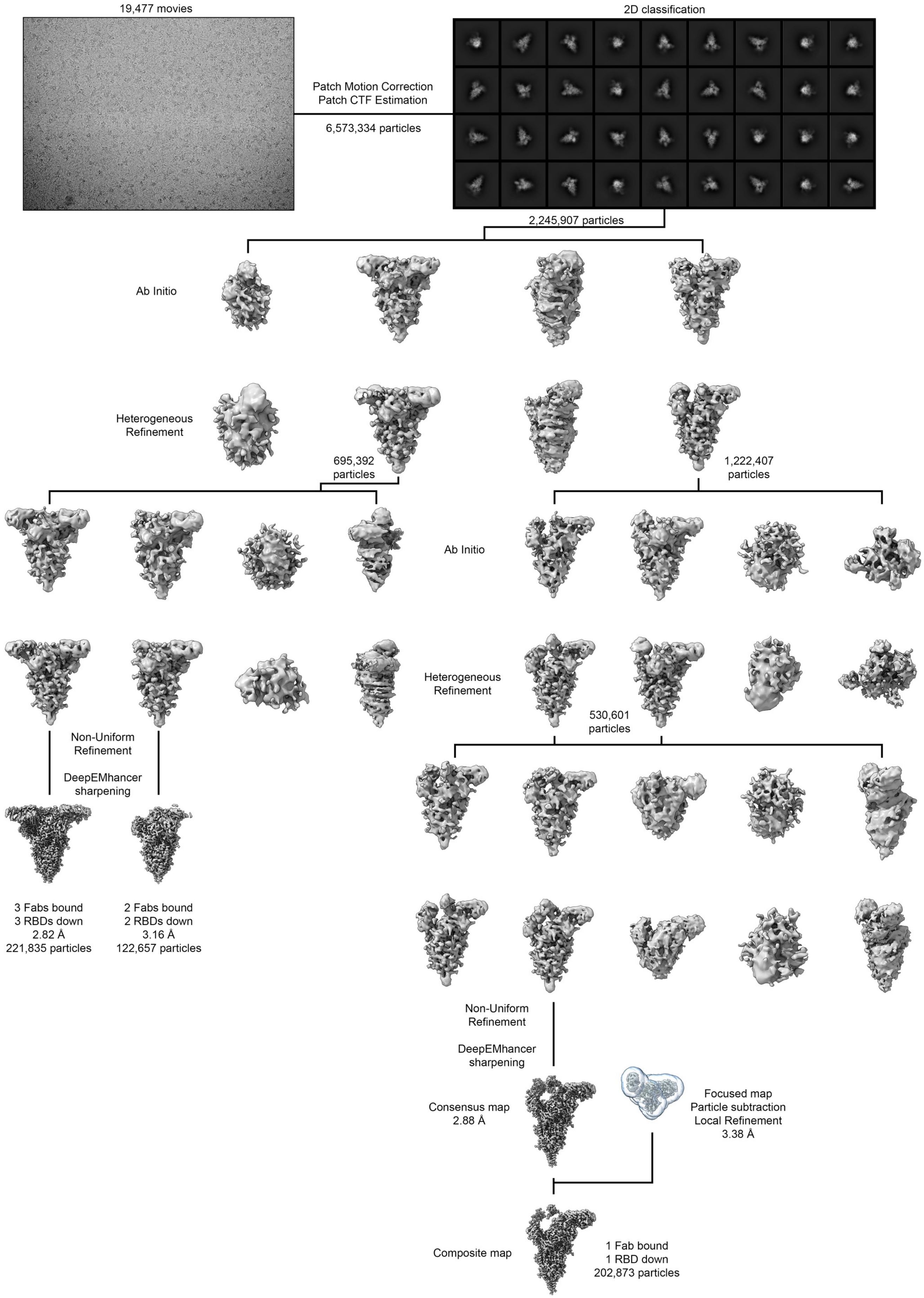


**Supplementary Figure 4: Cryo-EM data processing workflow.**

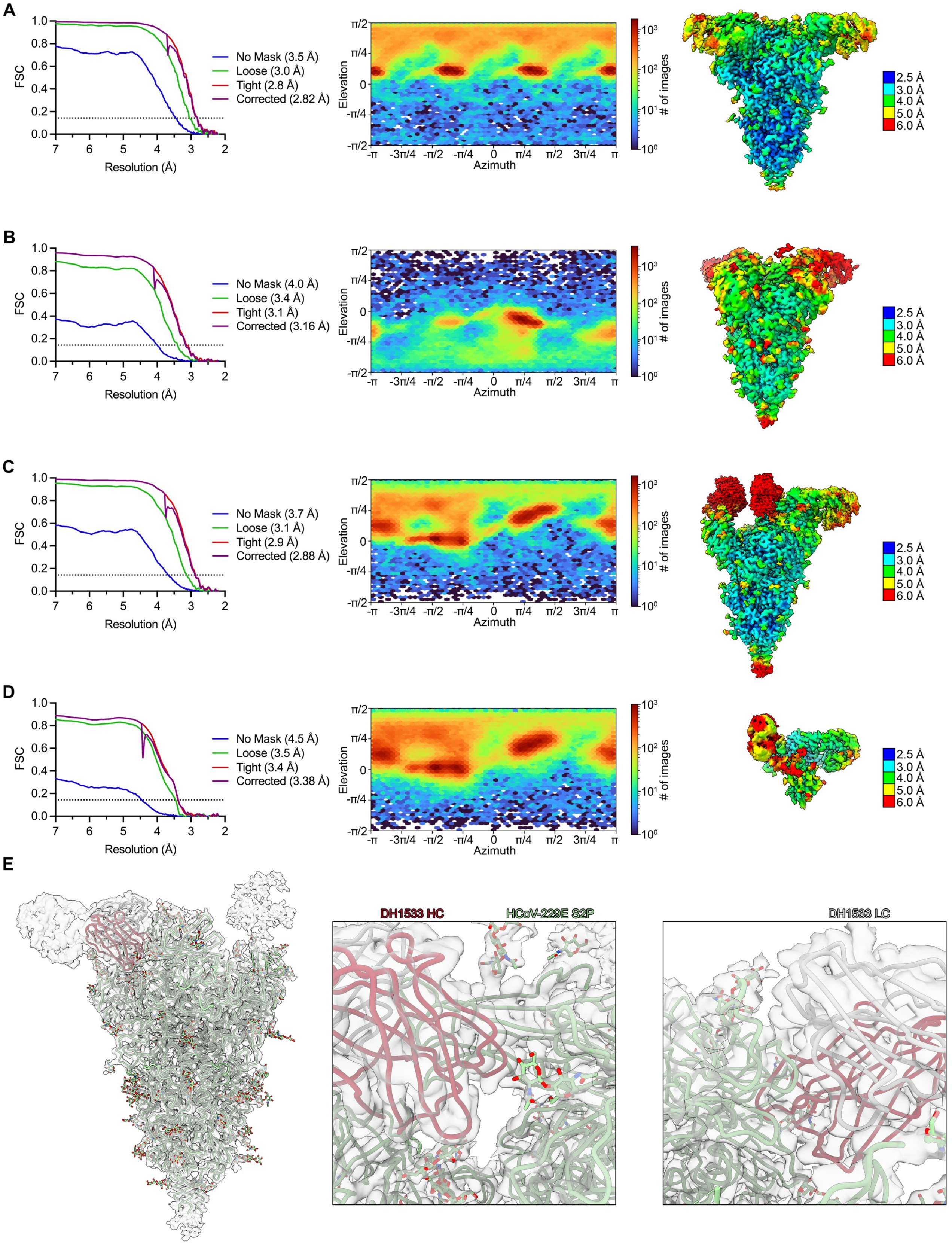


**Supplementary Figure 5: Cryo-EM validation**. The FSC curves (*left*), viewing direction distribution plots (*center*) and local resolution maps (*right*) are shown for (**A**) triply bound HCoV-229E S2P, (**B**) doubly bound HCoV-229E S2P, (**C**) singly bound HCoV-229E S2P, and (**D**) a local refinement of the singly bound complex, focused on an "up" RBD and the nearby DH1533-bound RBD. (**E**, *left*) The molecular model of HCoV-229E S2P bound by a single DH1533 Fab is shown docked into the composite map generated from the reconstructions shown in **C** and **D**. Zoomed-in views of the same model and map are shown, highlighting the DH1533 heavy chain (*left*) and light chain (*right*).

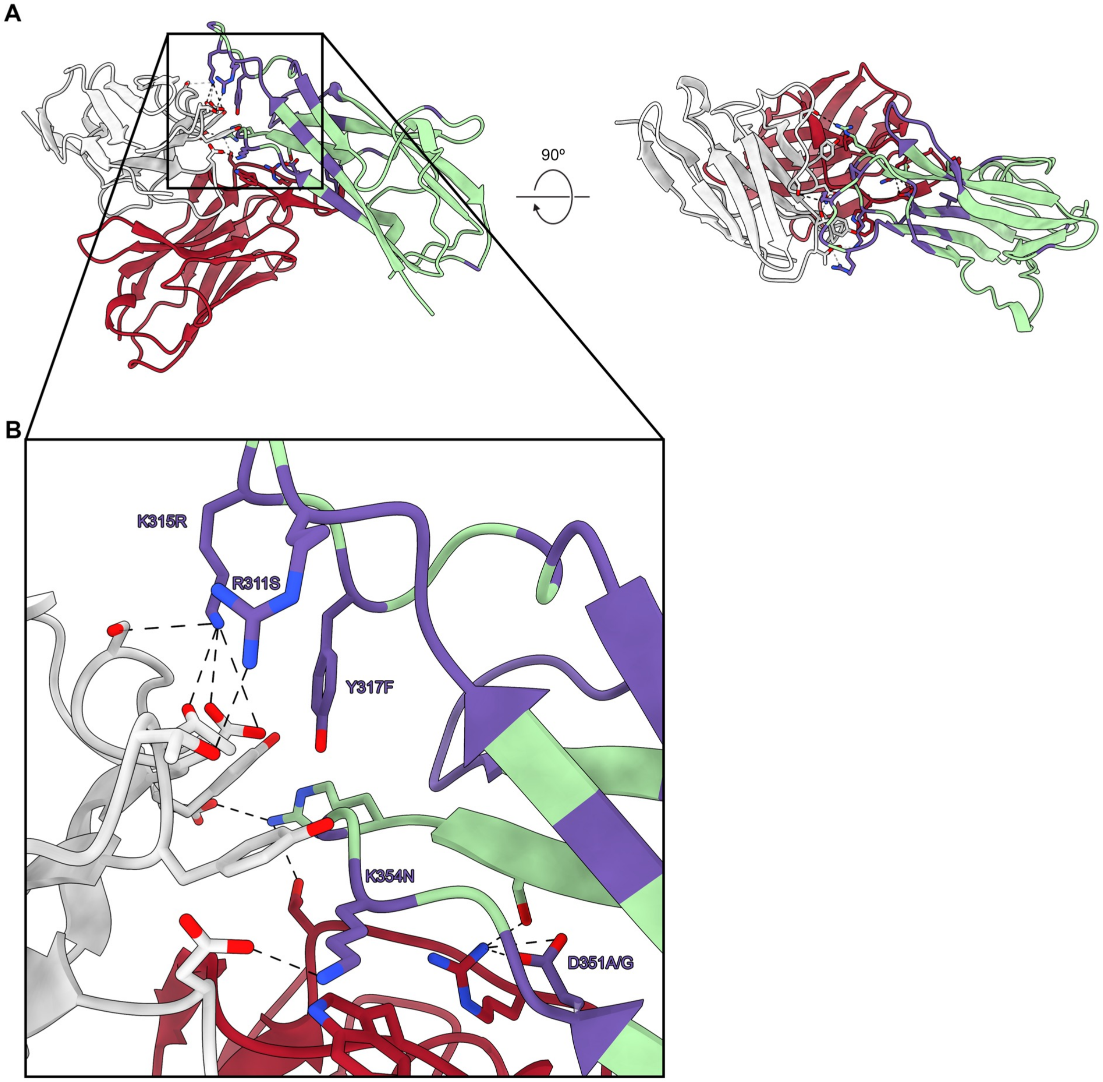


**Supplementary Figure 6: Structural analysis of antigenic drift in the DH1533 epitope**. (A) The structure of the variable domain of DH1533 bound to the RBD is shown from two different angles in ribbon diagrams, colored according to **Figure 4**. Residues that vary between the representative strains in **Supplementary Figure 3** are colored purple. (B) A zoomed-in view of the DH1533 binding interface. Critical contact residues are shown as sticks. Oxygen atoms are colored red, nitrogen atoms are colored blue and predicted hydrogen bonds are shown as black dashed lines. Critical contact residue mutations have been labeled.

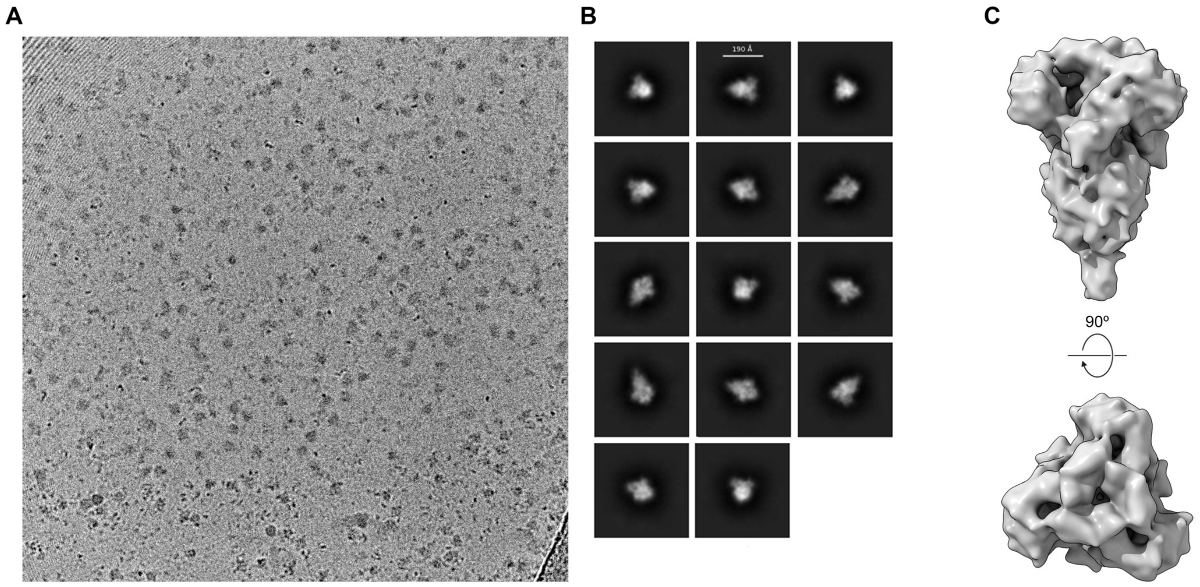


**Supplementary Figure 7: HCoV-229E S2P displays no conformational flexibility in the RBD in the absence of DH1533**. (**A**) Cryo-electron micrograph showing unbound HCoV-229E S2P trimers. (**B**) 2D class averages of HCoV-229E S2P particles. (**C**) An 8.1 Å 3D reconstruction of HCoV-229E S2P is shown from “side” and “top” views.

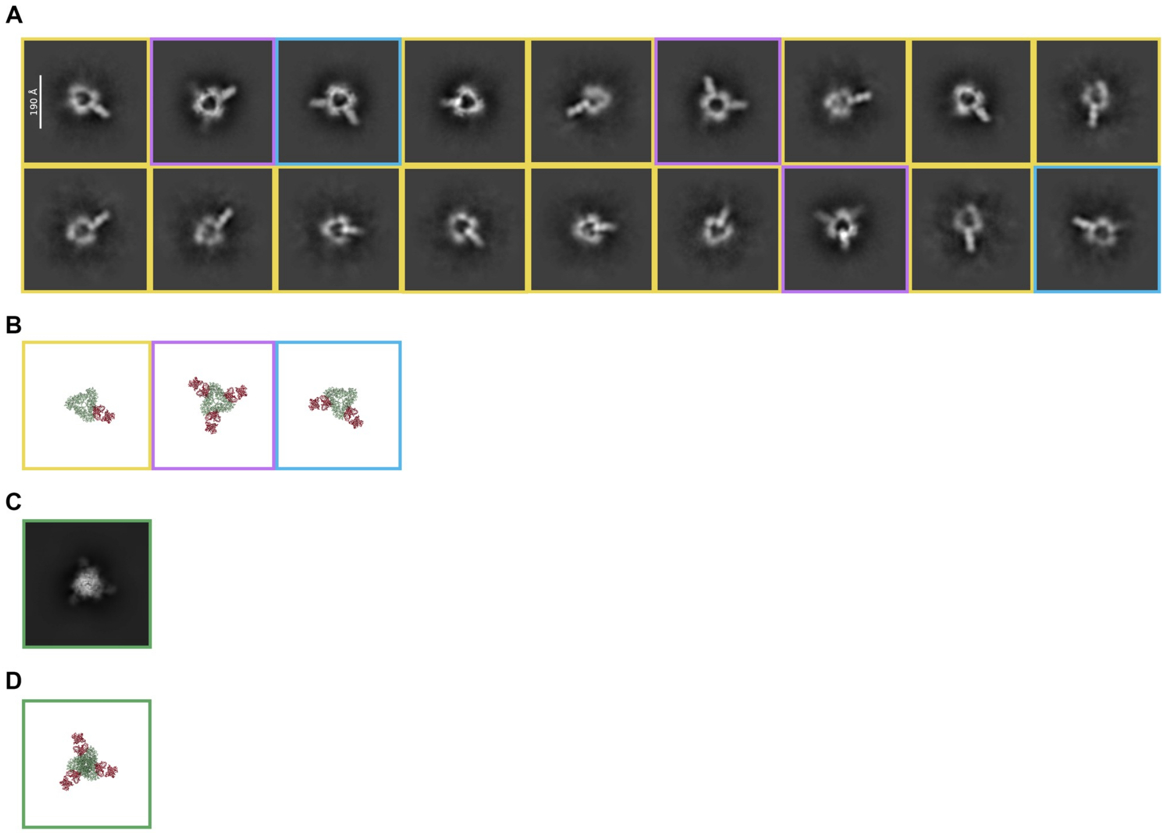


**Supplementary Figure 8: Dissociated S1 subunits observed by cryo-EM**. (**A**) 2D class averages from the DH1533 Fab + HCoV-229E S2P cryo-EM dataset that show dissociated S1 “rings”. Rings bound by a single DH1533 Fab have been highlighted yellow, rings bound by two DH1533 Fabs have been highlighted blue and rings with faint features suggesting three bound DH1533 Fabs have been highlighted purple. (**B**) Models corresponding to the 2D class averages in panel **A** are depicted, highlighted with the same coloring scheme. S1 ring models are colored green and DH1533 Fab models are colored red. (**C**) A 2D class average showing a “top” view of an intact, triply bound DH1533 Fab + HCoV-229E S2P complex has been highlighted green, with the corresponding model shown below in panel (**D**).

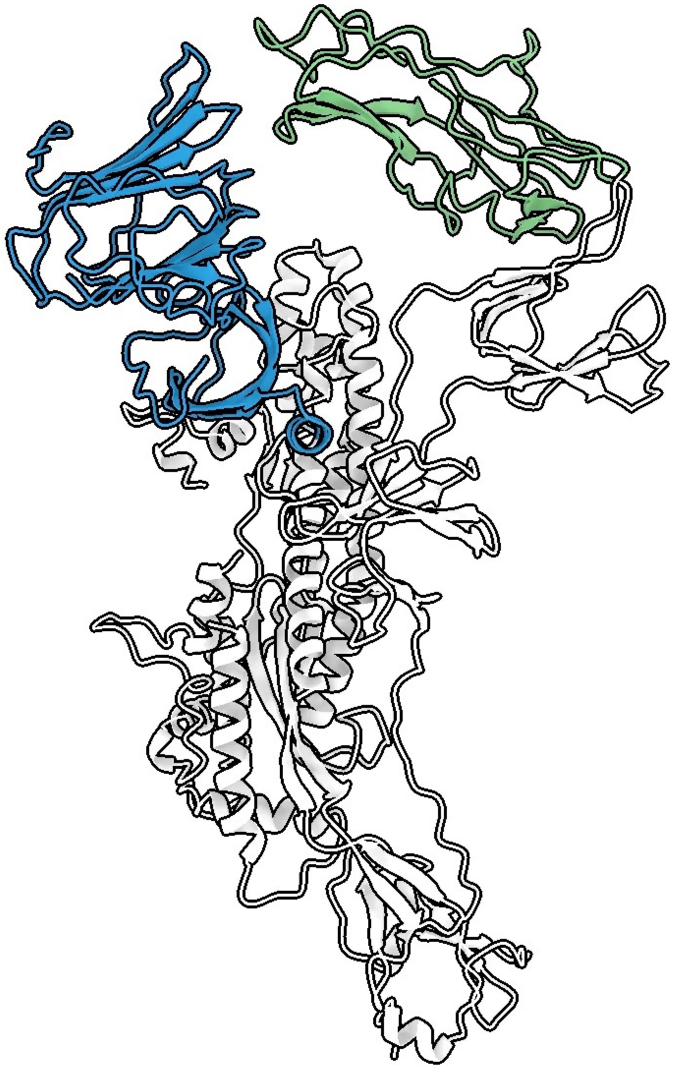

**Supplementary Movie 1: DH1533 binding causes compensatory movement in the neighboring NTD.** A single HCoV-229E S protomer is shown, interpolating the previously reported unbound conformation (PDB ID: 7CYC) and the DH1533-bound conformation. DH1533 binding causes the neighboring NTD to move downward, away from the RBD. The NTD is colored blue, the RBD is colored green and the rest of the protomer is colored white.

**cryo-EM Data Collection**

| | **HCoV-229E S2P + 3 DH1533 Fabs** | **HCoV-229E S2P + 2 DH1533 Fabs** | **HCoV-229E S2P + 1 DH1533 Fab** |
|---|---|---|---|
| Microscope | Titan Krios G3i | Titan Krios G3i | Titan Krios G3i |
| Voltage (kV) | 300 | 300 | 300 |
| Detector | K3 Bioquantum | K3 Bioquantum | K3 Bioquantum |
| Pixel size (Å/pix) | 1.08 | 1.08 | 1.08 |
| Exposure rate (e-/pix/sec) | 14.6 | 14.6 | 14.6 |
| Frames per exposure | 60 | 60 | 60 |
| Exposure ($e^-$/Å$^2$) | 49.8 | 49.8 | 49.8 |
| Defocus range (µm) | -0.8 – -2.0 | -0.8 – -2.0 | -0.8 – -2.0 |
| Micrographs used | 17,145 | 17,145 | 17,145 |
| Particles extracted/final | 6,573,334/221,835 | 6,573,334/122,657 | 6,573,334/202,873 |
| Symmetry | C3 | C1 | C1 |
| Resolution (Å) by FSC | | | |
| Unmasked 0.5 | 4.12 | 7.86 | 5.94 (8.06 local) |
| Masked 0.5 | 3.15 | 3.57 | 3.26 (3.93 local) |
| Unmasked 0.143 | 3.45 | 4.00 | 3.67 (4.45 local) |
| Masked 0.143 | 2.82 | 3.16 | 2.88 (3.38 local) |
| EMDB IDs | EMD-70440 | EMD-70441 | EMD-70442<br>EMD-70507<br>EMD-70508 |

**Model Refinement and Validation Statistics**

| | | | |
|---|---|---|---|
| Composition | | | |
| Amino acids | 3726 | 3357 | 2987 |
| Ligands | 126 | 104 | 171 |
| Bonds (RMSD) | | | |
| Length (Å) | 0.006 | 0.003 | 0.005 |
| Angles (°) | 1.34 | 0.89 | 1.02 |
| Ramachandran plot | | | |
| Outliers (%) | 0.1 | 0.2 | 0.2 |
| Allowed (%) | 3.5 | 4.7 | 4.1 |
| Favored (%) | 96.4 | 95.1 | 95.7 |
| Rotamer outliers (%) | 1.22 | 0.56 | 1.01 |
| Cβ outliers (%) | 0 | 0 | 0 |
| Clash score | 5.85 | 7.09 | 4.89 |
| MolProbity score | 1.61 | 1.73 | 1.56 |
| EMRinger score | 3.78 | 3.01 | 3.67 |
| PDB ID | 9OFO | 9OFP | 9OFQ |

**Supplementary Table 1: Cryo-EM data collection and refinement statistics.**
